# An anti-adhesive surface coating reduces adhesion during contact with cribellar threads in *Pholcus phalangioides* (Araneae, Pholcidae) but not in the web-owning spider *Uloborus plumipes*(Araneae, Uloboridae)

**DOI:** 10.1101/838250

**Authors:** Miriam Frutiger, Christian Kropf

## Abstract

Cribellar threads are powerful tools for web spiders to catch and retain prey. Spiders encountering such threads, like cribellar web spiders or araneophagic spiders invading cribellar webs, should have a protective mechanism against the adhesion of these threads. We tested for an anti-adhesive surface coating in the web invader *Pholcus phalangioides* and the cribellate orb weaver *Uloborus plumipes*. We calculated an index of adhesion for differently treated legs of the two species in a cribellar *U. plumipes* capture thread, i.e. untreated legs, water-washed legs, and legs washed with the organic solvent n-hexane. The results show that legs of *P. phalangioides* stick significantly stronger when washed with n-hexane. Our interpretation is that *P. phalangioides* has an organic surface coating lowering the adhesive force of the cribellar thread. No such mechanism was found in *U. plumipes*.

## Introduction

Many spiders construct webs and use adhesives to catch and retain prey. There are two mayor types of adhesion mechanisms used: threads equipped with droplets of viscous glue (1–3) and cribellar threads, composed of two axial threads surrounded by thousands of nano-fibrils (4,5). Any of these threads adhere strongly to a large variety of surfaces, but never to the web owning spider. This lack of adhesion is not purely due to the spiders behaviour, such as tiptoeing or avoidance of adhesive parts since it touches the capture spiral during construction as well as when struggling with large prey (2,6–8). So, the spider seems to have a protection against its own capture threads. Elaborate investigations in how this protection is formed were missing from scientific literature for many years. Two studies published in 2012 resumed the idea of the French naturalist Jean Henry Fabre from 1905, suggesting a putative lipid surface coating to prevent araneoid spiders of being caught in their gluey capture threads (6,8–10). By washing the legs of different araneoid spiders with either carbon disulphide or hexane, two different organic solvents dissolving lipids, they found an increased adhesive force between the spider’s leg and its own capture thread. It was concluded that these spiders either have chemical compounds on their epicuticle, which are washed off by the organic solvent, or some structural surface layer got altered by the chemicals applied (6,8).

Protective mechanisms against capture threads can be assumed in web-building spiders since their survival utterly depends on it. But what about spiders, that only occasionally enter such webs? Some spiders veered away from catching insects in their own webs only and further prey on other spiders or their eggs (11,12). A good example for versatility in predatory behaviour is the cellar spider *Pholcus phalangioides* (Fusselin, 1775) from the family Pholcidae: besides feeding on insects ensnared in its aerial sheet web, *P. phalangioides* also invades alien webs to feed on prey packages, egg sacks or the resident spider (12,13). While it hunts most successfully theridiid spiders in their own webs, it can also catch spiders in araneoid orbs or cribellate webs (13). A washable organic coating seems to protect *P. phalangioides* against the gluey capture threads of araneoid orb-weavers (14), but nothing is known about its protection against cribellar capture threads.

Cribellar capture threads operate with very different adhesive mechanisms as compared to gluey capture threads produced by araneoid spiders. They are dry composite threads, made up of two large axial threads and a mat of cribellar fibrils surrounding them (4,5) The cribellar fibrils act as adhesives, retaining insect prey even longer than gluey capture threads (15). The adhesion is achieved by interlock/entanglement, van der Waals and hygroscopic forces (16–18). A more recent study found yet another adhesive mechanism of the cribellar capture thread: the fibrils become embedded in the epicuticular waxes of the prey’s cuticula, increasing the adhesion by a multiple (19).Cribellar spiders, however, never get entangled in their own webs (M. F., personal observation). Studies about anti-adhesive mechanisms against cribellar capture threads in cribellar web spiders are absent.

This study investigates a possible protective mechanism against cribellar capture threads by focussing on two species that have either constant or temporary contact with cribellar capture threads: the cribellate orb-web spider *Uloborus plumipes* (Lucas, 1846) and the web-invader *Pholcus phalangioides* (13,20). We test both species for an anti-adhesive surface coating by washing their legs with n-hexane and measuring indirectly the force needed to detach differently treated spider legs from a cribellar thread spun by *U. plumipes*. We assume *P. phalangioides*, who uses gumfoot-lines in its own web (21), to have a putative wax coating, similar to araneoid spiders. Furthermore, we expect epicuticular waxes to interact with the cribellar fibrils and thus enhance adhesion as it was shown recently for the insect cuticle by Bott et al (2017). We therefore hypothesize that untreated legs of *P. phalangioides* adhere stronger to cribellar capture threads than those washed with an organic solvent (i.e. n-hexane), which should remove the wax layer. For the legs of *U. plumipes* we expect no difference in adhesion between the treatment groups because the adhesion mechanism of the cribellar thread as known yet does not favour a lipid surface coating as protection (19) but in contrast, this should even increase adhesion.

## Material and Methods

### Ethical statement

No endangered or protected species were used in this study. No special permits were required. All international, national and institutional guidelines for the care and use of animals were followed.

### Study animals and webs

#### Spiders

We collected *Pholcus phalangioides* in various households in the area of Bern, Switzerland, during the period of September-November 2018. Only large subadult individuals and adult females were used for measurements, males were excluded. The spiders were either used immediately after capture or kept in plastic tubes with a foam cone and a wetted tissue to keep moisture up to a maximum of 5 days.

*Uloborus plumipes* was collected in the Tropenhaus Frutigen, Switzerland, during September – December 2018. Here too, only large subadult individuals and adult females were used for measurements. The spiders were brought to the Natural History Museum Bern, Switzerland, and kept in plastic laboratory boxes (25*18*14 cm) where the spiders were allowed to spin their webs. The boxes were kept under warming lamps (24° Celsius), connected to a time-switch (12h light, 12h dark). The boxes were moistened directly before the spider was placed in the box and again after the web had been used for measurements. After a web was used for measurements, the spider was fed with fruit flies (*Drosophila melanogaster*) and provided with water.

#### Webs

For measurements we used webs of *U. plumipes*, spun in the laboratory boxes. The webs were 1-3 days old when used. 1 cm of the second outermost turn of the capture spiral of a web was cut out and fixed in a plastic object holder. From each web three threads were collected and randomly assigned to the three differently treated legs of a spider for later use. Legs of *U. plumipes* individuals were always tested in a web spun by another spider.

### Assignment and application of treatments

Spiders were euthanized in the freezer, then legs were cut off between femur and trochanter using forceps. In *P. phalangioides* legs I and II were used for measurements, in *U. plumipes* we used legs II and IV.

We applied 3 different treatments on the spider legs: 1) washing with n-hexane (**h**) for 2 minutes and drying it for 10 minutes under a fume hood, 2) leaving the leg untreated (**u**) but keeping it in a humidor for 12 minutes and 3) washing with double distilled water (**w**), then dry it for 10 minutes under a fume hood. The treatments were randomly assigned to each of the 3 legs dissected from one spider. Random samples were controlled under the stereomicroscope for complete drying. All measurements were conducted using a blind testing procedure, meaning that the measuring person had no knowledge on the respective treatment.

### Data collection

After application of treatments, a spider’s leg was fastened in a clamp (Fig. 1, a) on the net-o-meter 2.1 (Fig. 1, bamutec, bachofner museumstechnik gmbh, 3032 Hinterkappelen). An object holder containing a harvested thread was placed beneath the leg (Fig. 1, b). Then the leg was lowered, and the dorsal side of the tarsus was brought into touch with the thread until it applied a slight pressure on it (bending distance of the thread: 0.5mm). The control software (netometer V1.0.exe, developed by bamutec gmbh for the use of net-o-meter) was started, pulling the clamp with the leg upwards at a velocity of 20mm/s and an acceleration of 6m/s^2^. This speed was chosen because it lies within the range of velocities *U. plumipes* uses during thread production (7). A high-speed camera (Fig. 1, c; Optronis CR600X2-C-8G-GE-XM with a Schneider Kreuznach APO Componon 40/2.8 object lens, using TimeBench Software Ver. 2.6, Optronics GmbH) filmed the upward movement of the leg to the point where the leg detached from the thread, using a framerate of 500 frames per second. By counting the frames, we were able to calculate the exact distance at which the leg detached from the capture thread. This distance (in mm) we termed “Index of Adhesion” (IOA,(8)) and it served as an indirect measure of the force required to detach a spider’s leg from the capture thread.

**Fig 1:**
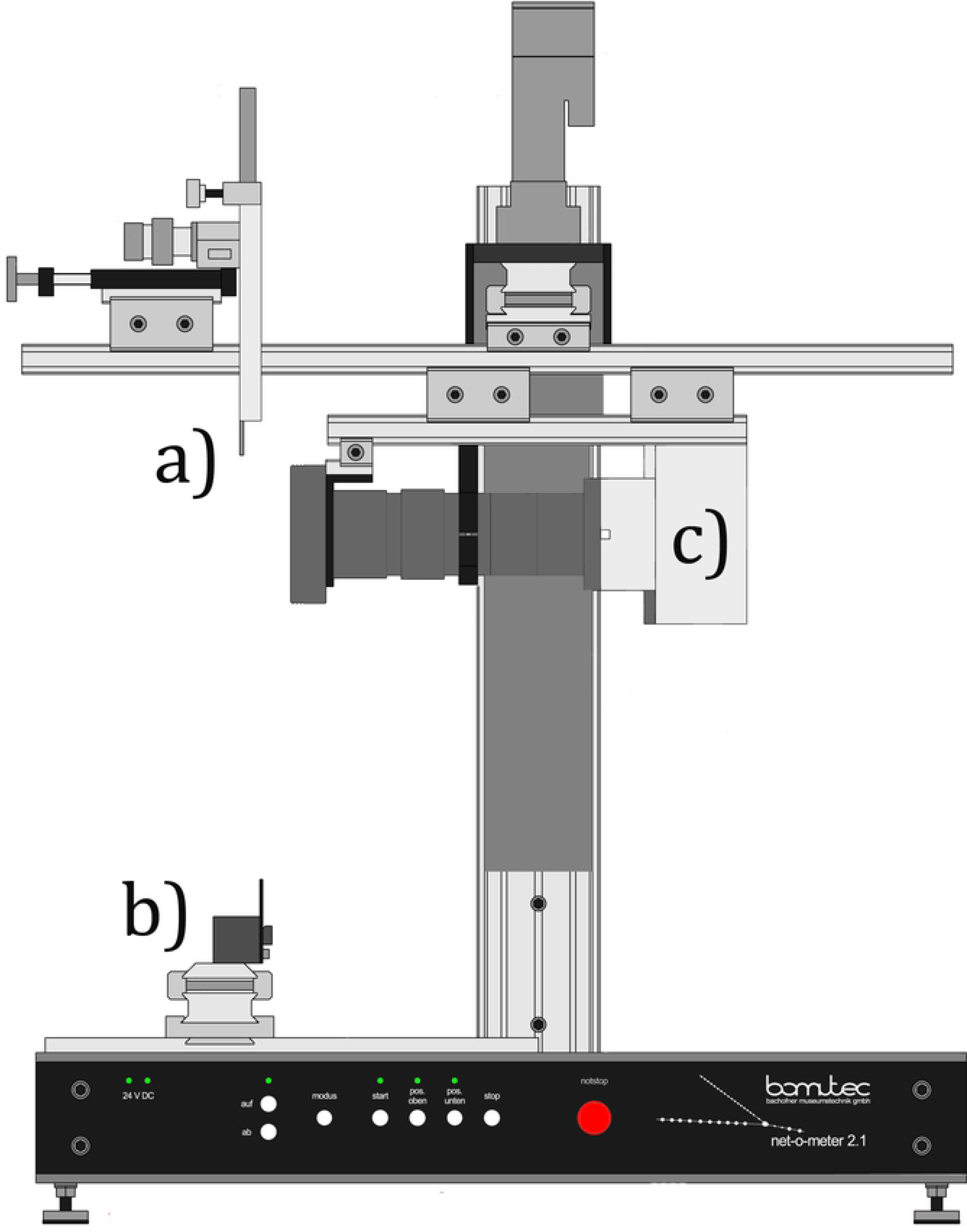
net-o-meter 2.1: a) clamp to fasten spider legs, b) moveable carrier to place threads on object holders, c) high-speed camera.

We used 80 specimens, 40 of each species, resulting in a total of 240 measurements, 40 measurements per treatment per species.

### Statistical Analysis

For the statistical analysis we compared the IOA of each treatment visually using boxplot in R (22). For the evaluation of the data we used a mixed-effects model with heteroscedastic residuals. This allowed us to identify tendencies in the mean IOA in treatments and species while correcting for the paired data points, arising from testing three legs per individual. These computations were performed using lme from the R package nlme (23). To test for differences in treatment effects within the species we performed post-hoc tests using lsmeans (24). All statistical computations and graphics were carried out in R software version 3.6.0 (25).

## Results

Our data clearly show an effect of washing on the legs of *P. phalangioides* (estimated means using lsmeans: u = 3.310 mm, w = 5.912 mm, h = 9.974 mm; see Fig. 2). The mean IOA value for legs washed with n-hexane is significantly higher than the one of untreated legs (p-value estimate using lsmeans = < .0001, SE = 0.205, df = 148), translating to a higher adhesive force between hexane-washed legs and cribellar thread compared to untreated legs. The water washing treatment shows values in between the untreated and hexane washed legs (see Fig. 2, p-value estimates u-w = 0.0148, SE = 0.205, df = 148; p-value estimates for h-w = 0.0314, SE = 0.205, df = 148).

**Fig 2:**
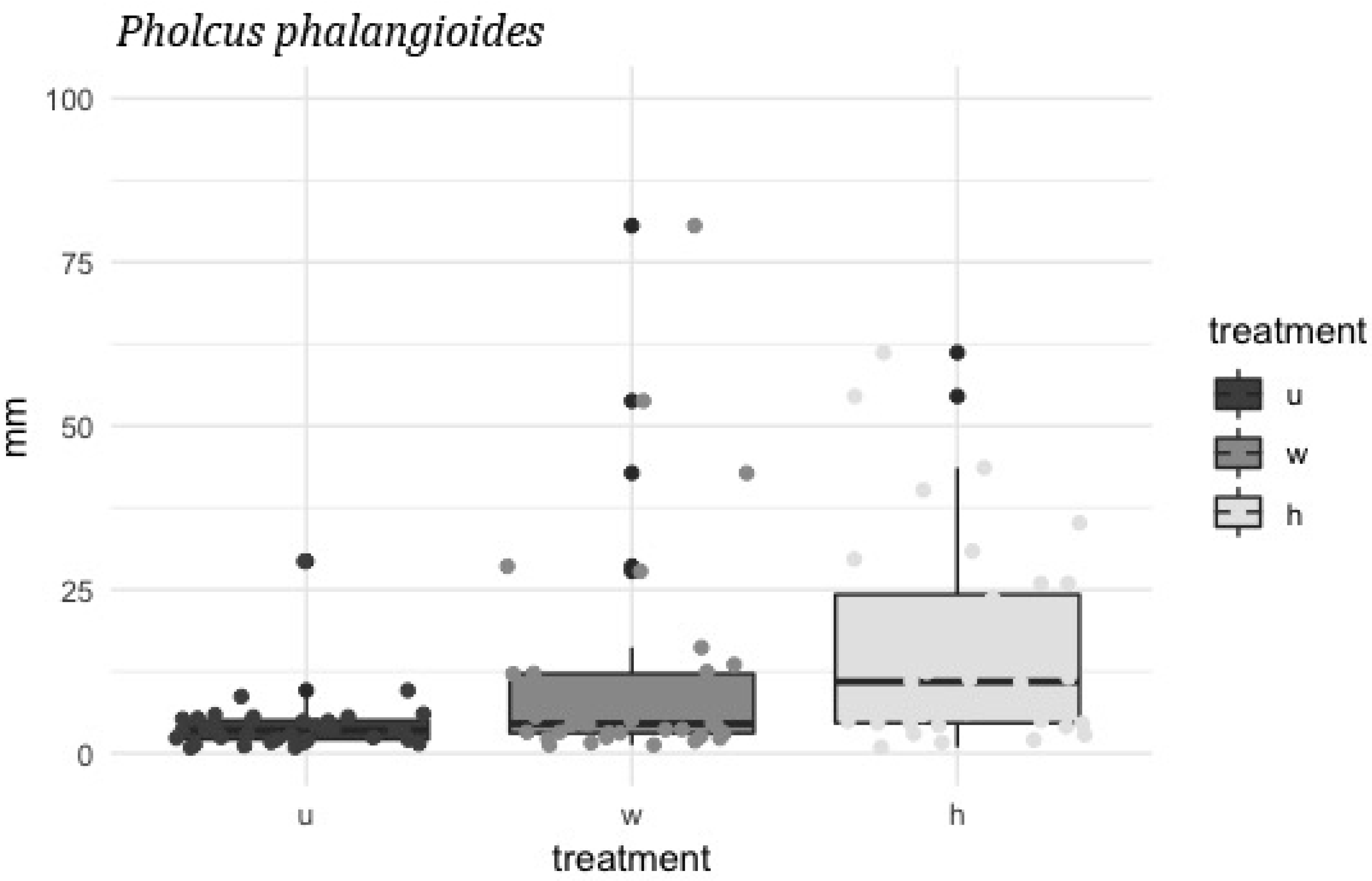
y-axe: detaching point of leg and thread in millimeters (IOA), x-axe: the different treatment groups in legs of *P. phalangioides:* h = hexane washed, u = untreated, w = water washed (n = 40 per group). The line through the box represents the median (second quartile), the lower part of the box the first and the top part the third quartile. The whiskers show the range Q1 – 1.5*IQR and Q3 + 1.5*IQR respectively while the black dots are outliers. The shaded dots represent the individual data points.

The measurements performed with legs of *U. plumipes* showed no effect of treatment at all (estimated means: u = 1.813 mm, w = 2.002 mm, h = 1.879 mm; see Fig. 3). The IOA values were generally low compared with the ones from *P. phalangioides*, and there is no significant difference between the treatment groups (p-value estimates: u-w = 0.748, u-h = 0.963, h-w = 0.887; SE = 0.136; df = 148). Comparing the IOA values of *P.phalangioides* to the values from *U.plumipes* using lsmeans post-hoc test, we found that legs of *P. phalangioides* adhere significantly stronger to cribellar threads, no matter what treatment was applied (p-value estimates for contrast between the species per treatment: u = 0.0019, w: = <.0001, h = <.0001; SE = 0.187; df = 74).

**Fig 3:**
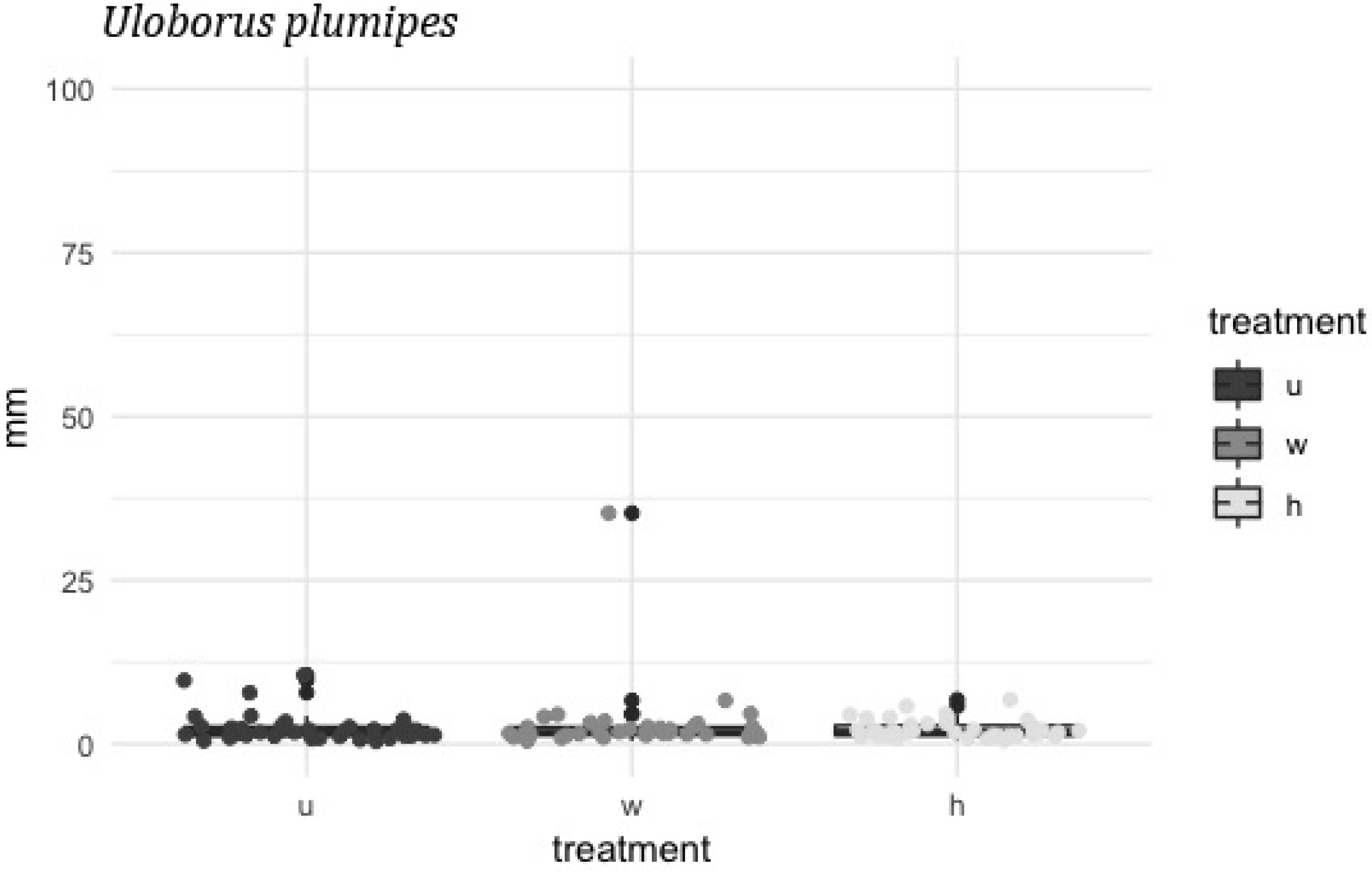
IOA in mm of the different treatment groups in *U. plumipes* legs; h = hexane washed, u = untreated, w = water washed (n = 40 per group).

## Discussion

Our data show that the IOA of *P. phalangioides* legs washed with n-hexane is increased by factor 3.01 when comparing to untreated legs. From this we conclude, that the legs surface has some kind of protection against cribellar capture threads that can be altered or dissolved by organic solvents. Former studies investigating the protective mechanism of web spiders obtained similar results. Kropf et al. (2012) found an increase of the IOA by factor 2.01 when comparing untreated and carbon disulphide-washed legs of *Araneus diadematus* on gluey capture threads spun by this species. Briceno and Eberhart (2012) likewise found a strong increase in force needed to break adhesion between the leg of *Nephila clavipes* and its gluey capture thread when the leg was washed with hexane. A direct comparison of the measurement values is not possible since all studies used different pull-off velocities, and these were shown to influence adhesion considerably (26).

Protection against viscous glue can either be achieved by chemically inert compounds or by microstructures minimizing the surface contact (27). An anecdotal study by Fabre (1905) states that a lubricated straw fails to adhere to the gluey capture web of araneoid spiders, pointing towards a chemical protection. Under this assumption, we expected different results when testing spider legs with the same methodology on gluey and cribellar capture threads: a lipid surface coating preventing adhesion on araneoid capture threads supposedly increases adhesion when coming in contact with the cribellar capture thread, because such surface lipids seemingly are soaked up by the cribellar nano-fibrils, in this way causing strong adhesion (19).To check if *P. phalangioides* has an epicuticular layer capable of interacting with the cribellar fibrils we examined cribellar capture threads contacting hexane-washed and untreated legs under the electron microscope. We found a considerable interaction in untreated legs, showing that the capture thread absorbs substances from the legs surface (Fig. 4 a). The cribellar fibrils obviously become embedded in some semi-liquid surface coating, changing the transparency of the thread (see (19)). In hexane-washed legs no such interaction was visible, indicating that the solvent actually washed off the semi-liquid waxes (Fig. 4 b).

**Fig 4:**
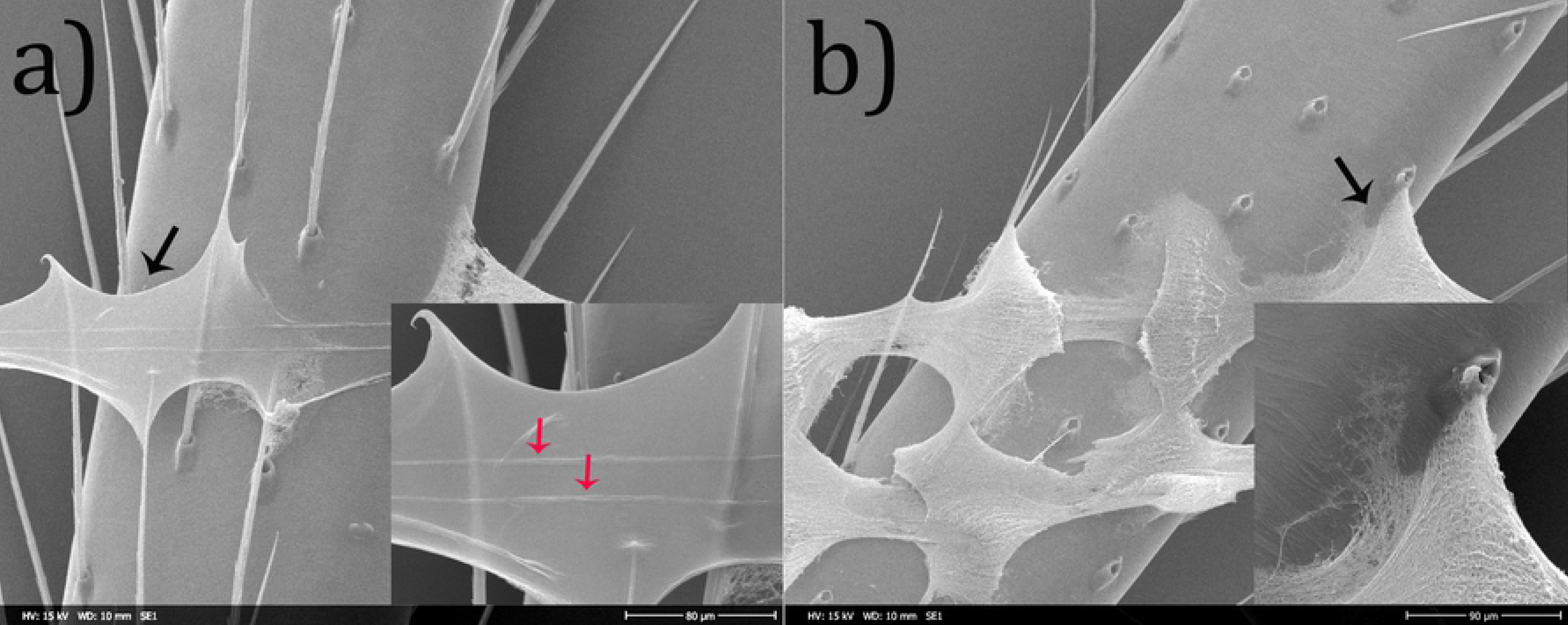
a) Untreated leg of *P. phalangioides*: the cribellar fibrils are embedded in the epicuticular waxes and form a semi-translucent sheet. Black arrow indicates position of insert, red arrows point out the axial threads. b) Leg of *P. phalangioides* washed with n-hexane: single fibrils are visible and probably adhering to the leg by van der Waals forces. Photo by A-C. Joel

A possible explanation for the decrease in IOA in untreated legs of P. phalangioides could lie in the fact that we measured distances and not the force needed for detachment. The epicuticular wax spreading in the cribellar fibrils could alter the mechanical properties of the capture thread, limiting a sensible comparison of the deflection of wax soaked threads vs native threads as indirect measurement of attachment forces (Joel, personal communication)., Legs of *P. phalangioides* washed with water show a mean IOA below the one of untreated and above the one of hexane-washed legs. Similar results were found by Briceno & Eberhard (2012), but an explanation is still pending.

The mean IOA of untreated *P. phalangioides’* legs exceeds the one of *U. plumipes* legs by factor 1.8. This indicates that, although the web invader seems to have a surface layer reducing adhesion, the protection against cribellar capture threads is still incomplete. This becomes visible when observing *P. phalangioides* walking in the web of *U. plumipes*: the invader cannot move freely in a cribellar orb web but gets more or less stuck on the capture thread (video of a *P. phalangioides* moving through the orb web of *U. plumipes* can be accessed here: https://doi.org/10.6084/m9.figshare.10028339.v1). It is though capable of freeing itself, probably partly due to the protective surface coating as well as to its long legs, enabling it to reach a non-sticking surface and drag itself out of the web. Extensive grooming of the legs and biting through of cribellar threads are also behaviours observed in *P. phalangioides*, which are highly probable of facilitating its performance in the cribellar web (13).

Finding reduced mobility of *P. phalangioides* in webs of *U. plumipes* puts up the question of how the web invader preys on cribellar spiders. In our laboratory as well as in greenhouses, where both species co-occur, we observed, that the *Pholcus*’ web often is woven adjacent to cribellar webs, even appear to be fused, with *P. phalangioides* adjusting its own silk lines on the cribellar web. This might suggest, that *P. phalangioides* does not enter the cribellate web completely but rather builds bridges of his own silk. Jackson and Brassington (1987) observed *P. phalangioides* to perform a vibratory behaviour when entering a cribellate web, which often resulted in the resident spider approaching the web invader. To learn more about the hunting behaviour of *P. phalangioides* in cribellate webs and the frequency with which it chooses to invade such webs, more behavioural studies would be needed.

Measuring the adhesion of legs of *U. plumipes’* on cribellar capture threads, we found no difference in IOA between the treatment groups and continuously low adhesion. This suggests, that no organic coating altering the adhesive force between tarsus and capture threads is present in this spider. Legs of cribellate spiders show a microscale pattern of little grooves on their cuticle while ecribellate spiders mostly show a pattern resembling scales (28,29). Microscale patterns on the exo-and epicuticle of arthropods have been suggested before to have anti-adhesive properties (30,31). Perhaps the protective mechanism of cribellate spiders against their own capture threads can be found in these cuticular structures.

Our results pose more questions on how the cribellar thread interacts with epicuticular substances. So far only preliminary results regarding the chemical composition of the epicuticular waxes of web spiders are accessible (6). For further investigations in the anti-adhesive surface coating of *P. phalangioides* extensive knowledge about the composition and origin of surface lipids in web spiders is needed.

## Aknowledgements

We thank Wolfgang Nentwig and Marina Marthaler for making the data collection of this study possible, Anna-Christin Joel for helpful discussions and suggestions, Christof Strähl and Anja Mühlemann for statistical support /services. MF thanks Karin Urfer and Jonathan Berger for unfailing support.

